# Discovery and biosynthetic assessment of *Streptomyces ortus* sp nov. isolated from a deep-sea sponge

**DOI:** 10.1101/2022.11.21.517041

**Authors:** Sam E. Williams, Catherine R. Back, Eleanor Best, Judith Mantell, James E. M. Stach, Tom A. Williams, Paul R. Race, Paul Curnow

## Abstract

The deep sea is known to host novel bacteria with the potential to produce a diverse array of undiscovered natural products. Understanding these bacteria is thus of broad interest in ecology and could also underpin applied drug discovery, specifically in the area of antimicrobials. Here, we isolate a new strain of *Streptomyces* from the tissue of the deep-sea sponge *Polymastia corticata* collected at a depth of 1869 m from the Gramberg seamount in the Atlantic Ocean. This strain, which was given the initial designation A15ISP2-DRY2^T^, has a genome size of 9.29 Mb with a GC content of 70.83%. Phylogenomics determined that A15ISP2-DRY2^T^ represents a novel species within the genus *Streptomyces* as part of the *Streptomyces aurantiacus* clade. The biosynthetic potential of A15ISP2-DRY2^T^ was assessed relative to other members of the *aurantiacus* clade via comparative gene cluster family (GCF) analysis. This revealed a clear congruent relationship between phylogeny and GCF content. A15ISP2-DRY2^T^ contains six unique GCFs absent elsewhere in the clade. Culture-based assays were used to demonstrate the antibacterial activity of A15ISP2-DRY2^T^ against two drug-resistant human pathogens. We thus determine A15ISP2-DRY2^T^ to be a novel bacterial species with considerable biosynthetic potential and propose the systematic name *Streptomyces ortus* sp. nov.

**Impact Statement:** The *Streptomyces* genus has contributed more to our antibiotic arsenal than any other group of bacteria or fungi. Despite decades of exploration, global analysis has suggested they still possess more undiscovered biosynthetic diversity than any other bacterial group. Isolating novel species of *Streptomyces* is therefore a priority for antibiotic discovery. Here we isolate a novel strain from a deep-sea sponge and use comparative cluster analysis to identify six biosynthetic clusters unique to our deep-sea strain. This work demonstrates the utility of continuing to isolate novel *Streptomyces* strains for antibiotic discovery and, for the first time, we used species tree-gene cluster tree reconciliation to assess the contribution of vertical evolution on the biosynthetic gene cluster content of *Streptomyces*.

## Introduction

Antimicrobial resistance (AMR) is a major threat to human health and was associated with nearly 5 million deaths worldwide in 2019 (1). The discovery of new antibiotics with novel modes of action is a critical part of combatting this threat. In recent years there has been a renewed focus on microbial natural products as the basis for this discovery (2). Historically, the majority of antibiotic natural products have come from the bacterial group actinomycetes, with the majority of these arising from a select few genera such as *Streptomyces* and *Micromonospora* (3, 4). It has been estimated that over 50% of all clinically used antibiotics are derived from the Streptomycetes alone (4-7).

While *Streptomyces* remain an attractive target for biodiscovery initiatives, these efforts can be generally frustrated by the continual re-discovery of known natural products (8). One way to mitigate this problem is to focus upon relatively under sampled environmental niches that could harbour strains which have acquired biosynthetic innovations (9, 10). The microbial fauna intimately associated with marine sponges have long been seen as a potential source of novel bioactive metabolites, and *Streptomyces* strains are known to feature in sponge microbiota (11, 12). While the bioprospecting of sponge samples has largely been limited to accessible shallow waters (13), a more ‘extreme’ niche occupied by sponges – the deep ocean – has been less well explored (14). It is now apparent that deep-sea sponges can indeed host microbial communities with impressive bioactivity, and that many such communities include the Streptomycetes (15-17).

As well as traditional culture-based methods, bioinformatic genome mining for biosynthetic gene clusters (BGCs) now has a central role in the process of natural product discovery (5, 18). The identification of biosynthetic clusters by tools such as antiSMASH (19) is complemented by software that can, for example, predict whether a particular gene cluster might produce an antibiotic (20). Grouping together similar BGCs from multiple genomes into Gene Cluster Families (GCFs) has provided a deeper understanding of global biosynthetic diversity (21-23) and has been used to estimate that just 3% of bacterial natural product classes have so far been discovered (24). In such analyses, *Streptomyces* are found to have the greatest biosynthetic potential with certain phylogenetic subgroupings within the genus – specifically, a clade termed ‘*Streptomyces*_RG1’ (Relative Evolutionary Distance or RED group 1) – expected to be biosynthetically exceptional even by the standards of other *Streptomyces* (24). The isolation, identification, and characterisation of novel taxa within this ‘RG1’ clade is therefore a priority for natural product discovery.

Here, we describe the discovery of a novel species of deep-sea *Streptomyces* with a diverse biosynthetic repertoire and inherent antibiotic activity. We investigate the biosynthetic potential of this strain through a comparative analysis of GCF diversity within the *Streptomyces aurantiacus* clade (25) and highlight the extent that specialised metabolism within this clade remains unexplored.

## Results

### Isolation of Strain A15ISP2-DRY2^T^

A bacterial strain given the in-house designation A15ISP2-DRY2^T^ was isolated on ISP2 agar as part of an ongoing effort to culture bacteria from the microbiota of deep-sea sponges (26). The sponge sample was originally recovered by remote-operated vehicle from the Gramberg seamount in the Atlantic Ocean (depth 1869m; latitude 15.44530167; longitude: -51.09606). This was identified as the demosponge *Polymastia corticata* based on cytochrome oxidase I (COI) gene identity (Table S1, GenBank: OP036683). Strain A15ISP-DRY2^T^ produced a zone of inhibition against the Gram-positive test strain *Bacillus subtilis* in a soft agar overlay assay but did not inhibit the growth of the Gram-negative test strain *Escherichia coli* (data not shown). A15ISP2-DRY2 ^T^ was shown to be Gram-positive (Figure S1), and electron microscopy of the strain revealed branching filamentous substrate and spores (Figure 1).

**Figure 1.**
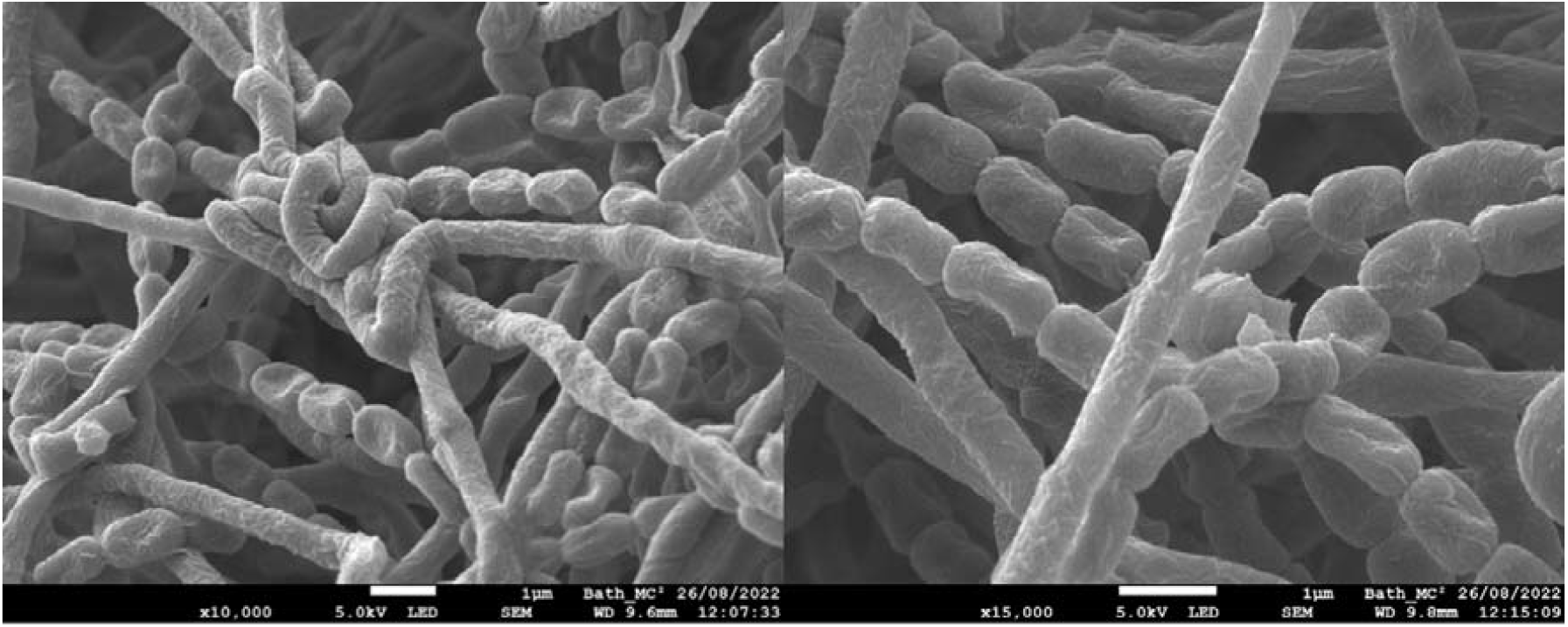
Cryo-Scanning Electron Micrographs of A15ISP2-DRY2 ^T^ colony grown on Mannitol Soya Flour Medium for 4 days.

### Taxonomic assignment

A full-length 16S rRNA gene sequence of 1525 bp (GenBank: ON356025.1) was sequenced and submitted to NCBI BLASTN 2.13.0+. A maximum likelihood tree of closely-related type strains indicated that the deep-sea isolate was a member of the *Streptomyces aurantiacus* clade (25) (Figure 2). Within this clade the closest relatives to the isolate were the strains *Streptomyces dioscori* A217^T^ and *Streptomyces liliiviolaceus* BH-SS-21^T^ with 16S rRNA gene identities of 99.74% and 99.61%, respectively.

**Figure 2.**
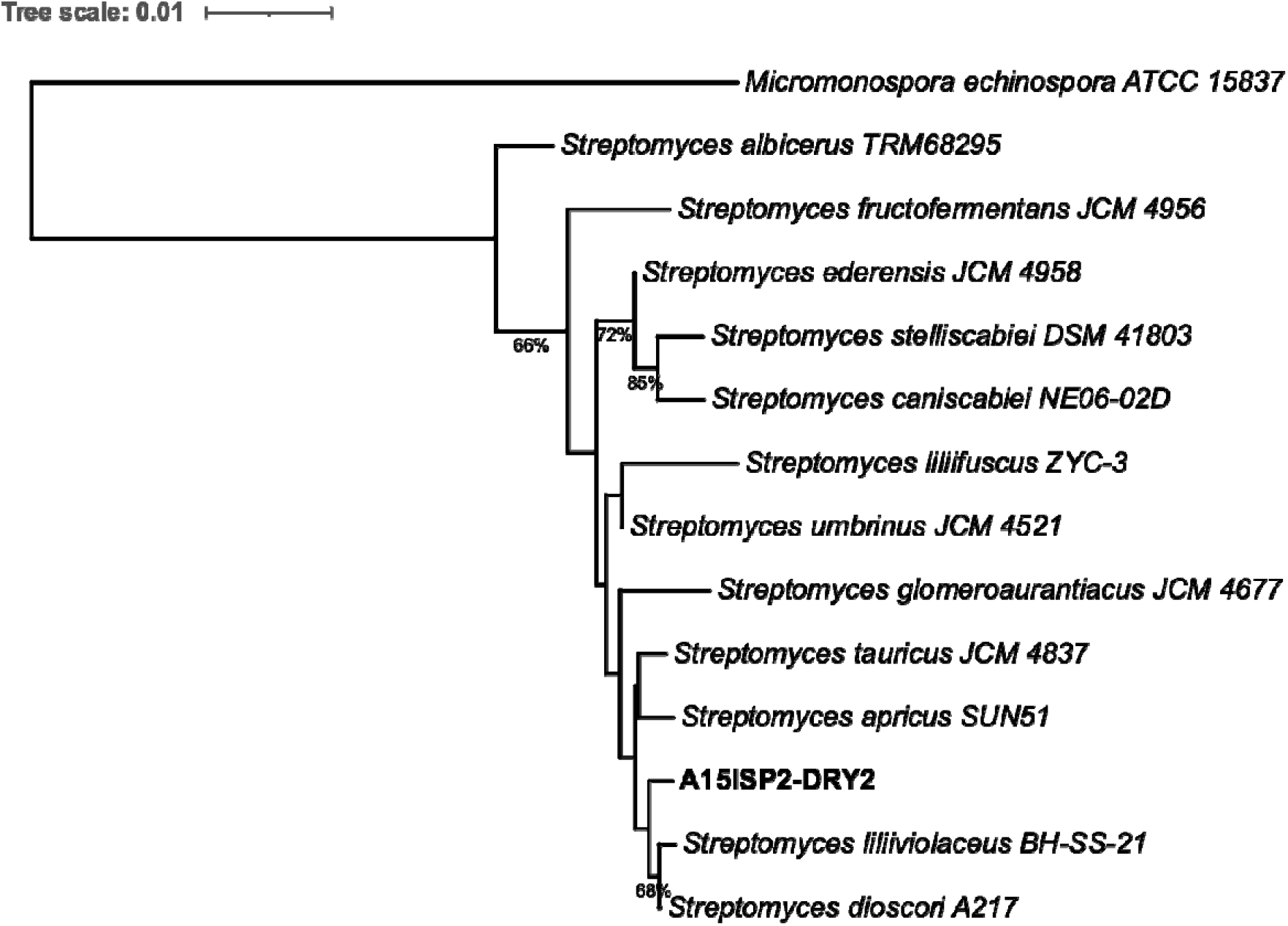
16S rRNA gene sequence phylogeny of A15ISP-DRY2^T^. Produced with The Type (Strain) Genome Server (TYGS), bootstrap values below 50% not shown, average branch support 49.5%, 14 strains, δ statistics = 0.455. Micromonospora echinospora ATCC 15837 is used as an outgroup and the tree is rooted on this branch.

The differential chemotaxonomic properties of strain A15ISP2-DRY2^T^ and the closely related *Streptomyces* species *S. dioscori* JCM 32173^T^, *S. liliviolaceus* BH-SS-21^T^ and *S. tauricus* JCM 4837^T^ are given in Table 1. Major fatty acids (>10%) for strain A15ISP2-DRY2^T^ were anteiso-C_15:0_, C_16:0_, iso-C_16:0_ and C_16:1_ω7*c*. Minor fatty acids (>5%) were iso-C_14:0_, iso-C_15:0_ and anteiso-C_17:0_. Whole cell sugars detected were glucose, galactose, ribose, and minor amounts of mannose. The diagnostic amino acid in whole-cell hydrolysates was LL-2,6-diaminopimelic acid. Menaquinones MK8 H6 10.5%, MK8 H8 0.8%, MK9 H4 8.1%, MK9 H6 50.5% and MK9 H8 30.1% were present.

**Table 1.**
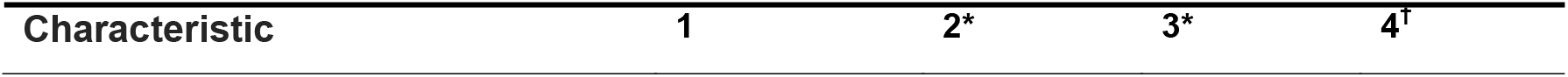
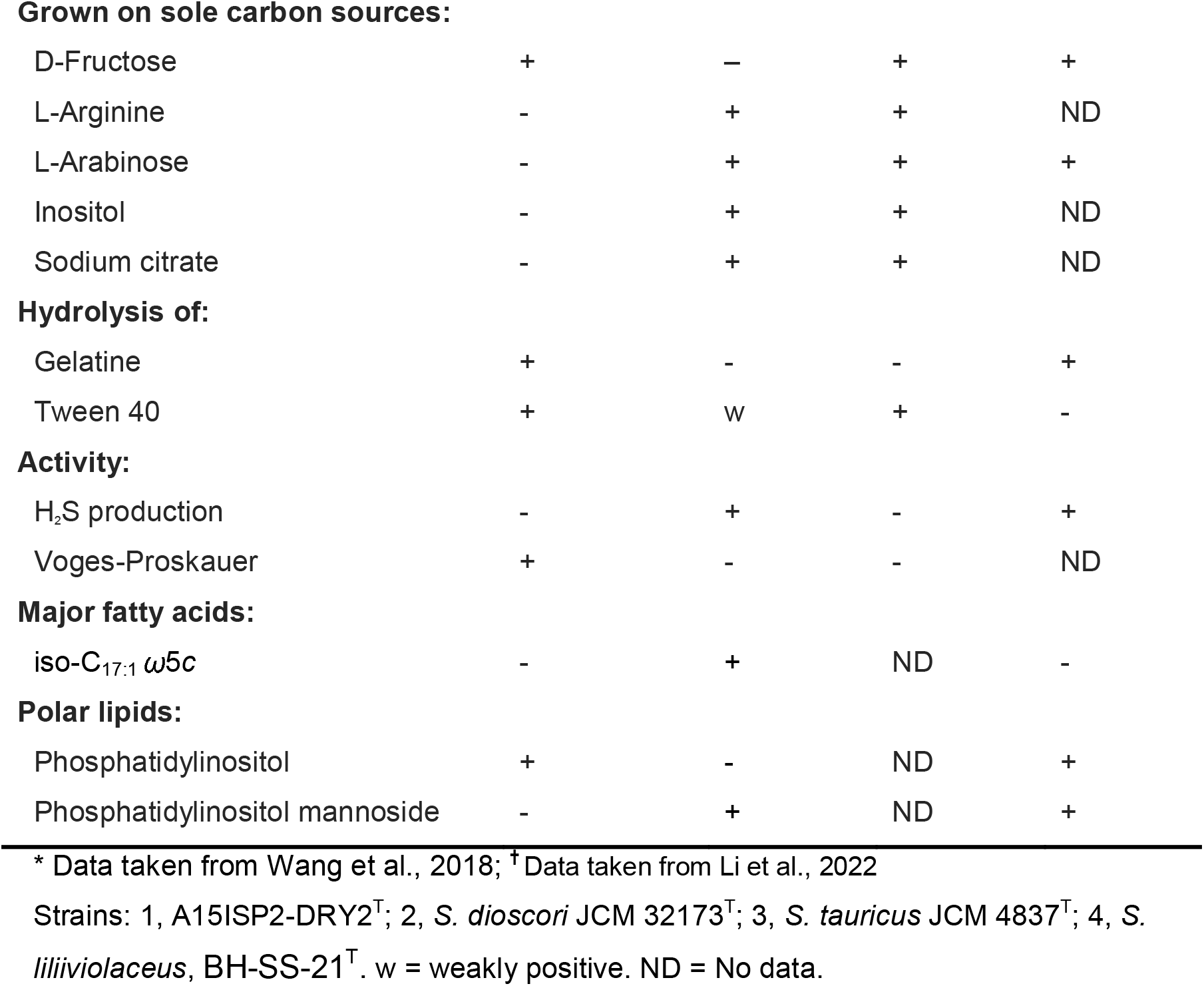
Differential characteristics of strain A15ISP2-DRY2^T^ and closely related Streptomyces type strains

| Characteristic | 1 | 2* | 3* | 4<sup>†</sup> |
| --- | --- | --- | --- | --- |
| **Grown on sole carbon sources:** |  |  |  |  |
| D-Fructose | + | — | + | + |
| L-Arginine | - | + | + | ND |
| L-Arabinose | - | + | + | + |
| Inositol | - | + | + | ND |
| Sodium citrate | - | + | + | ND |
| **Hydrolysis of:** |  |  |  |  |
| Gelatine | + | - | - | + |
| Tween 40 | + | w | + | - |
| **Activity:** |  |  |  |  |
| H<sub>2</sub>S production | - | + | - | + |
| Voges-Proskauer | + | - | - | ND |
| **Major fatty acids:** |  |  |  |  |
| iso-C<sub>17:1</sub> ω5c | - | + | ND | - |
| **Polar lipids:** |  |  |  |  |
| Phosphatidylinositol | + | - | ND | + |
| Phosphatidylinositol mannoside | - | + | ND | + |

\* Data taken from Wang et al., 2018; † Data taken from Li et al., 2022
Strains: 1, A15ISP2-DRY2<sup>T</sup>; 2, *S. dioscori* JCM 32173<sup>T</sup>; 3, *S. tauricus* JCM 4837<sup>T</sup>; 4, *S. liliiviolaceus*, BH-SS-21<sup>T</sup>. w = weakly positive. ND = No data.

The genome of A15ISP2-DRY2^T^ was sequenced to allow complete taxonomic assignment and the biosynthetic potential of this strain. The complete assembled genome was 9,291,524 bp in length, with a GC content of 70.83 %. The assembly consisted of 9 contigs, or 4 scaffolds, with an L50 of 1 and the largest scaffold being 8,605,295 bp in length (Table S2). The assembled genome contained 8,130 genes, six complete rRNAs, 66 tRNAs and a single CRISPR array (Table S3). The genome was further analysed for expected single-copy orthologous genes from the order Streptomycetales. Of 1579 expected genes, 1574 were complete (99.7%), suggesting the assembly was of high biological accuracy (Table S4).

The Type (Strain) Genome Server (https://tygs.dsmz.de) (27) was used to calculate digital DNA-DNA hybridisation (dDDH) values and to create a whole-genome phylogeny (Figure 3). FastANI was then used to report the average nucleotide identity (ANI) between the isolated strain and closely related type strains (28) (Table S5). This was consistent with the results of 16S analysis and confirmed that the closest relatives to A15ISP2-DRY2^T^ are *S. liliiviolaceus* BH-SS-21^T^, with dDDH score of 45.8%, ANI 93.31% and %GC difference 0.04, and ‘*S. dioscori* A217^T^’ with dDDH 45.1%, ANI 93.08% and %GC difference 0.11. The relative dissimilarity of the dDDH and ANI scores between A15ISP2-DRY2^T^ and these related strains reveals that A15ISP2-DRY2^T^ should be considered a distinct species. We thus propose here that A15ISP2-DRY2^T^ be given the systematic name *Streptomyces ortus* sp. nov. A species description, including an explanation of the epithet and details of culture deposition in accessible collections, is provided at the end of this manuscript.

**Figure 3.**
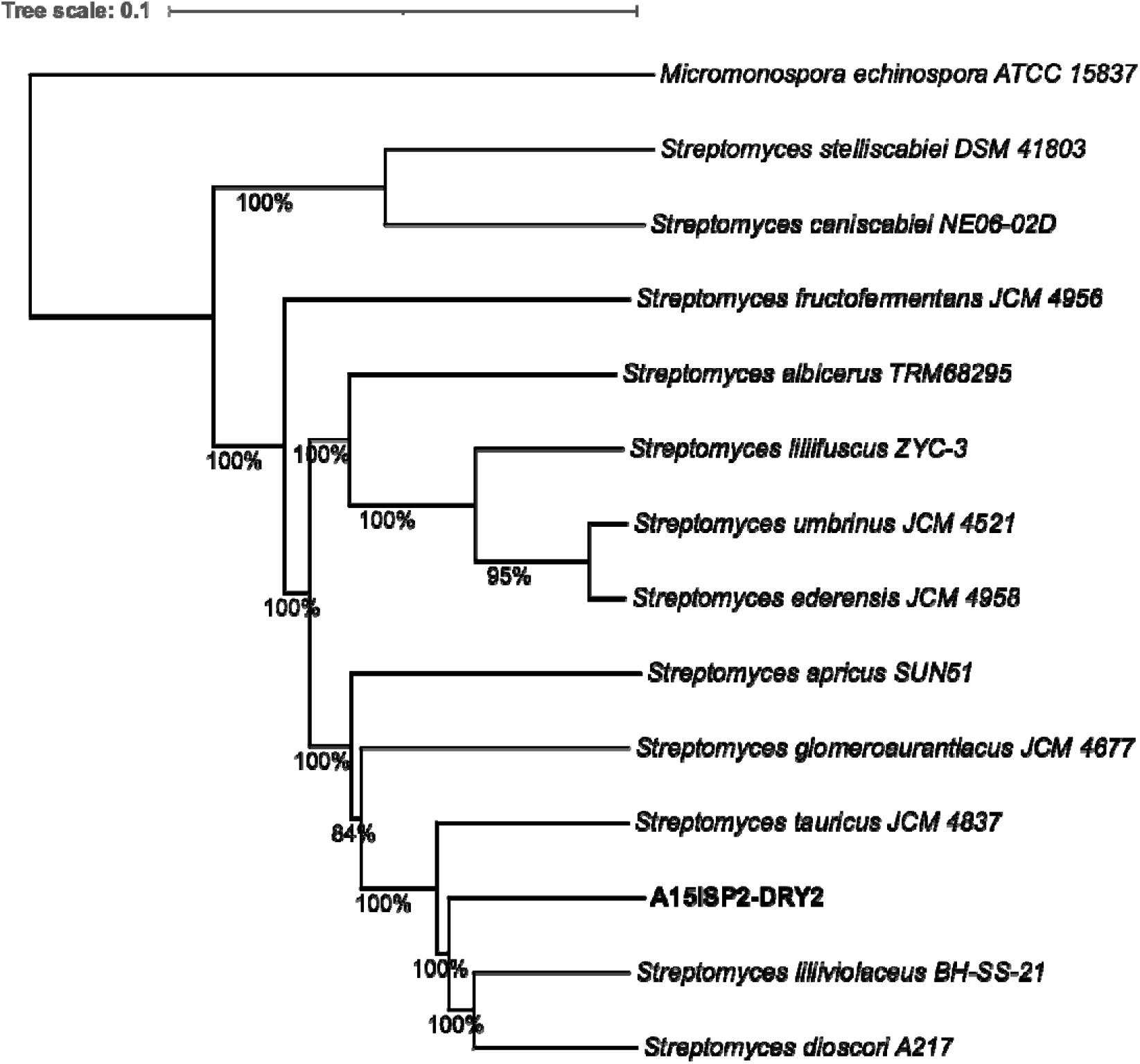
Maximum likelihood whole-genome phylogeny. 13 strains automatically chosen using the Type Genome Server (TYGS) with GGDC. Micromonospora echinospora ATCC 15837 is included as an outgroup and the tree is rooted on this branch. Average branch support 98.1%, bootstrap data shown as % for each branch. δ statistics = 0.131. Excluding the outgroup: % GC 70.02-72.41, genome size 7.67-11.96 Mb, Number of. proteins 6,307-10,784. Distance formula = D5

The whole-genome analysis additionally revealed that A15ISP2-DRY2^T^ belongs to the recently defined *Streptomyces*_RG1, the clade which has the highest biosynthetic potential of any genus level group in the bacterial kingdom (24). This suggests that the *Streptomyces* isolate described here represents an excellent candidate for further bioprospecting.

### Analysis of biosynthetic gene clusters

The genome of A15ISP2-DRY2^T^ was analysed with antiSMASH 6.1.1 for the identification of putative biosynthetic gene clusters (BGCs). A total of 34 complete BGCs were identified. Just 9 of these showed high gene similarity (>80%) to currently characterised BGCs (Table 2). Only 19 of the identified BGCs were most similar to those found in the closest relative listed in the antiSMASH database ‘*S. dioscori* A217’ (Table S6). Additionally, the genome was submitted to the Antibiotic Resistant Target Seeker 2 (ARTS; https://arts.ziemertlab.com) (29). This identified that 19 of the 34 identified BGCs in A15ISP2-DRY2^T^ were in proximity to a duplicated core gene or a known antibiotic resistance gene. This genomic context indicates that these BGCs may produce antibiotic compounds (18, 30).

**Table 2.**
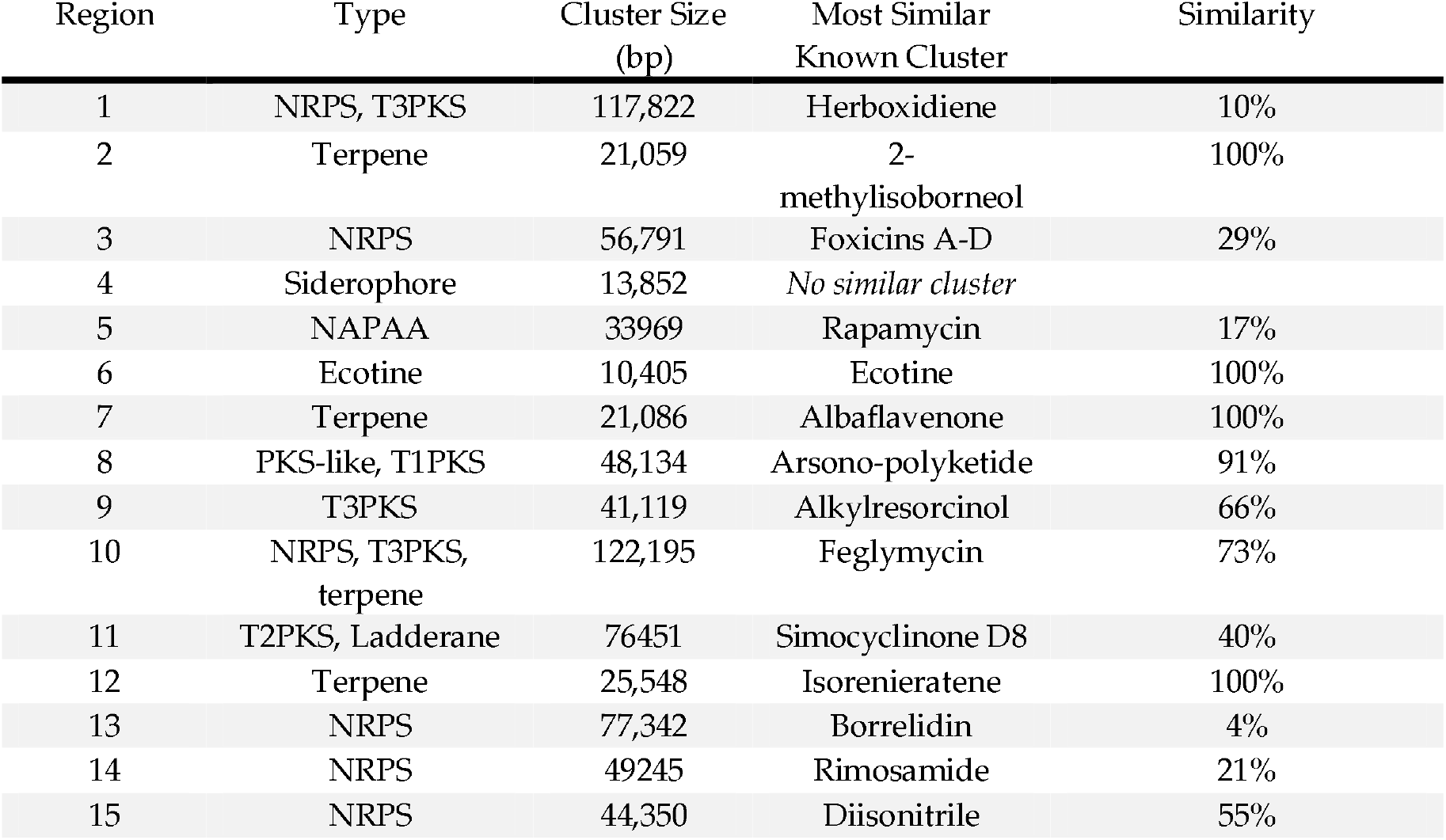
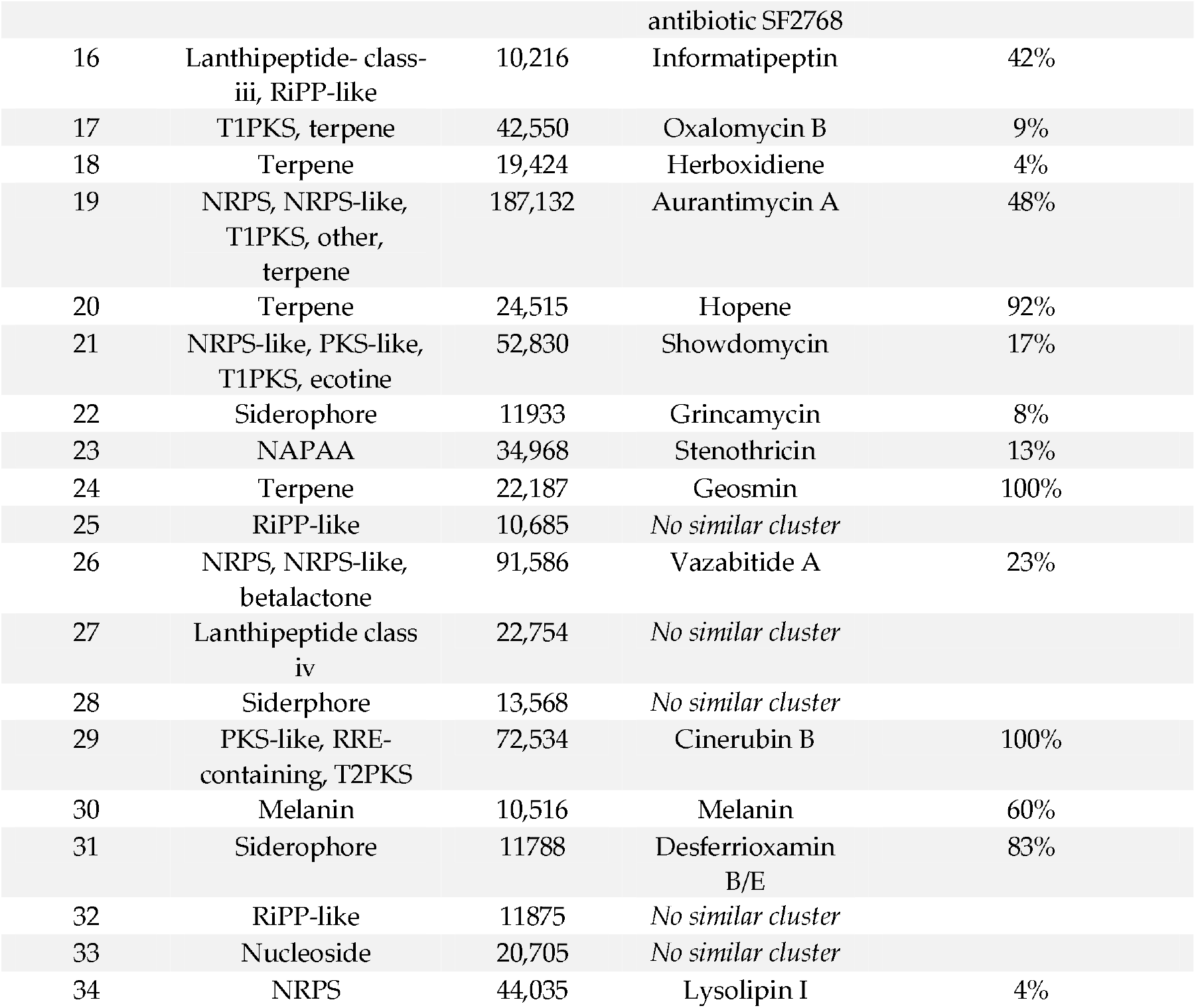
Output from antiSMASH 6.1.1 for the genome assembly of isolate A15ISP2-DRY2^T^.

### Biosynthetic Cluster Comparison within the *S. aurantiacus* Clade

To further evaluate the biosynthetic novelty of A15ISP2-DRY2^T^, a BiG-SCAPE gene cluster family (GCF) analysis (21) was conducted within a clade of the 10 closest relatives. A total of 377 BGCs were identified across the 10 genomes analysed. These grouped into 188 GCFs with 133 singletons and a total of 577 links (Figure 4, Figure S2). Just 22 (11.7%) of these GCFs clustered with a MiBIG reference BGC (31), highlighting the extent of unexplored specialised metabolism within this clade. Just six biosynthetic GCFs were shared amongst all members of the clade (Figure 4). Less stringently, ten GCFs were near-ubiquitous, being conserved in all members of the clade apart from *S. glomeroaurantiacus* JCM 4677^T^. The six conserved GCFs included the well-characterised clusters responsible for producing Ectoine, Hopene, Geosmin, Desferrioxamin B/E and Alkylresorcinol which are common across most *Streptomyces* strains (32). The other GCF found ubiquitously across the clade was a siderophore cluster with low gene similarity (8%) to that for the antibiotic polyketide Grincamycin (33).

**Figure 4.**
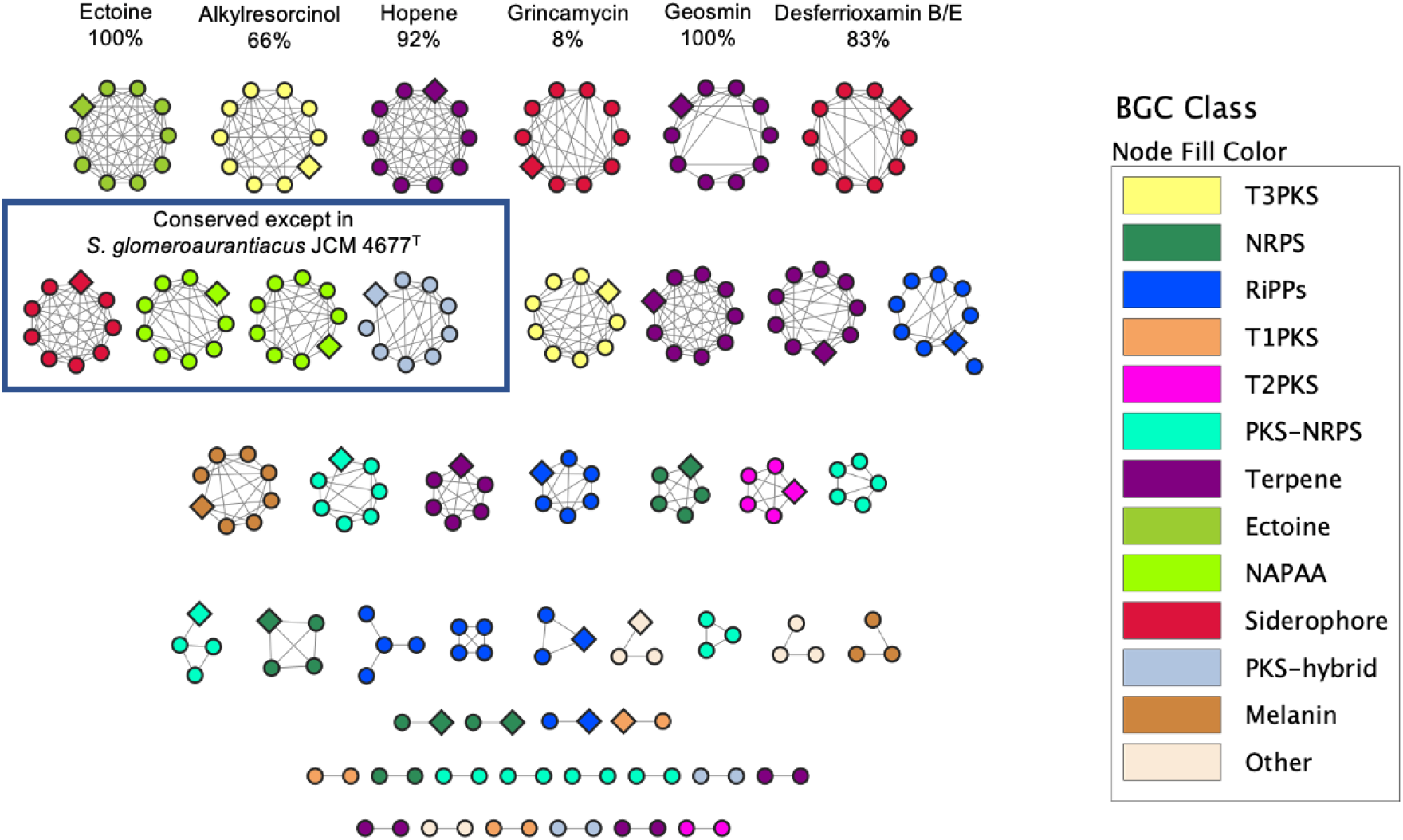
BiG-SCAPE GCF network analysis of 10 Streptomyces within the S. auranticus clade, including A15ISP2-DRY2^T^. Conserved GCFs are annotated with the name of the compound produced by the most similar known cluster, and antiSMASH % gene similarity to that cluster. Diamonds nodes are BGCs belonging to A15ISP2-DRY2^T^. Fragmented BGC nodes were removed so that only one node per GCF per strain was left in the network. Singletons are excluded from the figure. Figure legend shows compounds produced by gene products of the most similar known cluster. Key: NRPS: non-ribosomal polyketide synthase, here including NRPS-like clusters; RiPP, ribosomally-synthesised and post-translationally modified peptides, here including lanthipeptides and RiPP-like clusters; NAPAA: non-alpha poly-amino acids; PKS-hybrid, here including T1PKS-terpene, heterocyst glycolipid synthase-like PKS-T1PKS, Other, here including redox-cofactor, phosphoglycolipid and nucleoside. T1PKS: Type I polyketide synthase, T2PKS: Type II polyketide synthase, T3PKS: Type III polyketide synthase.

The phylogenetic subgroup encompassing A15ISP2-DRY2^T^, *S. liliiviolacens* BH-SS-21^T^, *‘S. dioscori’* A217 and *S. tauricus* JCM 4837^T^ exclusively contained GCFs for a polyketide synthase-non-ribosomal polyketide synthase (PKS-NRPS) hybrid cluster and an NRPS cluster. The subgroup also contained the type II polyketide synthase (T2PKS) for the anthracycline cinerubin B (34). Interestingly the cinerubin B GCF was also found in *S. fructofermentans*, suggesting this GCF has moved into the clade on two separate events. A nucleoside GCF with no similar other known cluster was found in all members of the group except *S. tauricus*. A15ISP2-DRY2^T^ and its closest relative *S. liliiviolacens* BH-SS-21^T^ also had two GCFs not found elsewhere including a NRPS-like cluster with 23% similarity to a cluster for Vazabitide A (35). This BGC was deemed likely to produce an antimicrobial, with ARTS 2.0 identifying a duplicated core gene and a putative resistance model within the cluster. Overall, this analysis demonstrates a clear phylogenetic relationship of BGC distribution in this clade, with close relatives sharing a higher portion of conserved BGCs.

A15ISP2-DRY2^T^ had 6 GCFs not found in any other members of the clade, potentially suggesting 6 BGC acquisition events since its speciation from *S. liliiviolacens*. Two of these unique GCFs are likely to produce antibiotics based on ARTS analysis. These are a T2PKS and ladderane hybrid BGC, containing a known resistance model and 40% gene homology to the novel DNA gyrase inhibitor simocyclinone D8 (36); and an NRPS BGC with a duplicated cell envelope core gene within the cluster. A15ISP2-DRY2^T^ also had a unique large NRPS-type III polyketide synthase (T3PKS) hybrid BGC with 73% similarity to the glycopeptide-related antibiotic feglymycin, but ARTS did not detect any resistance markers for this cluster. There thus appear to be uncharacterised BGCs within A15ISP2-DRY2^T^, responsible for producing antibiotics and not found in closely related *Streptomyces* species.

### Predominantly Vertical GCF Transmission within the *S. aurantiacus* Clade

To investigate the contributions of vertical inheritance and lateral gene transfer (LGT) to the evolution of BGCs within the *S. aurantiacus* clade, we performed gene cluster tree-species tree reconciliation. In total there were 28 GCF trees, where the GCF was present in at least 3 strains, which were reconciled against the clade species tree using an Amalgamated Likelihood Estimation (ALE) (37). ALE draws the gene tree into the species tree using a probabilistic model of gene origination, duplication, transfer, and loss, with model parameters estimated from the data using maximum likelihood. Consistent with the observed congruence between the species tree and BGC repertoires of these organisms, the ALE analysis suggested that vertical inheritance was the predominant mode of BGC evolution within the *S. aurantiacus* clade. The GCF branch verticality, a measure of vertical transmission in relation to transfer or loss, demonstrated an average GCF verticality of 89%. This analysis suggested that 12 to 13 of the GCFs analysed were already present in the last common ancestor of the *S. aurantiacus* clade (Figure 5). The analysis also highlighted loss events, such as a T1PKS-NRPS BGC with homology to aurantimycin A, and evidenced acquisition events, such as a T1PKS-NRPS BGC with low homology to showdomycin in the *S. ortus* group.

**Figure 5.**
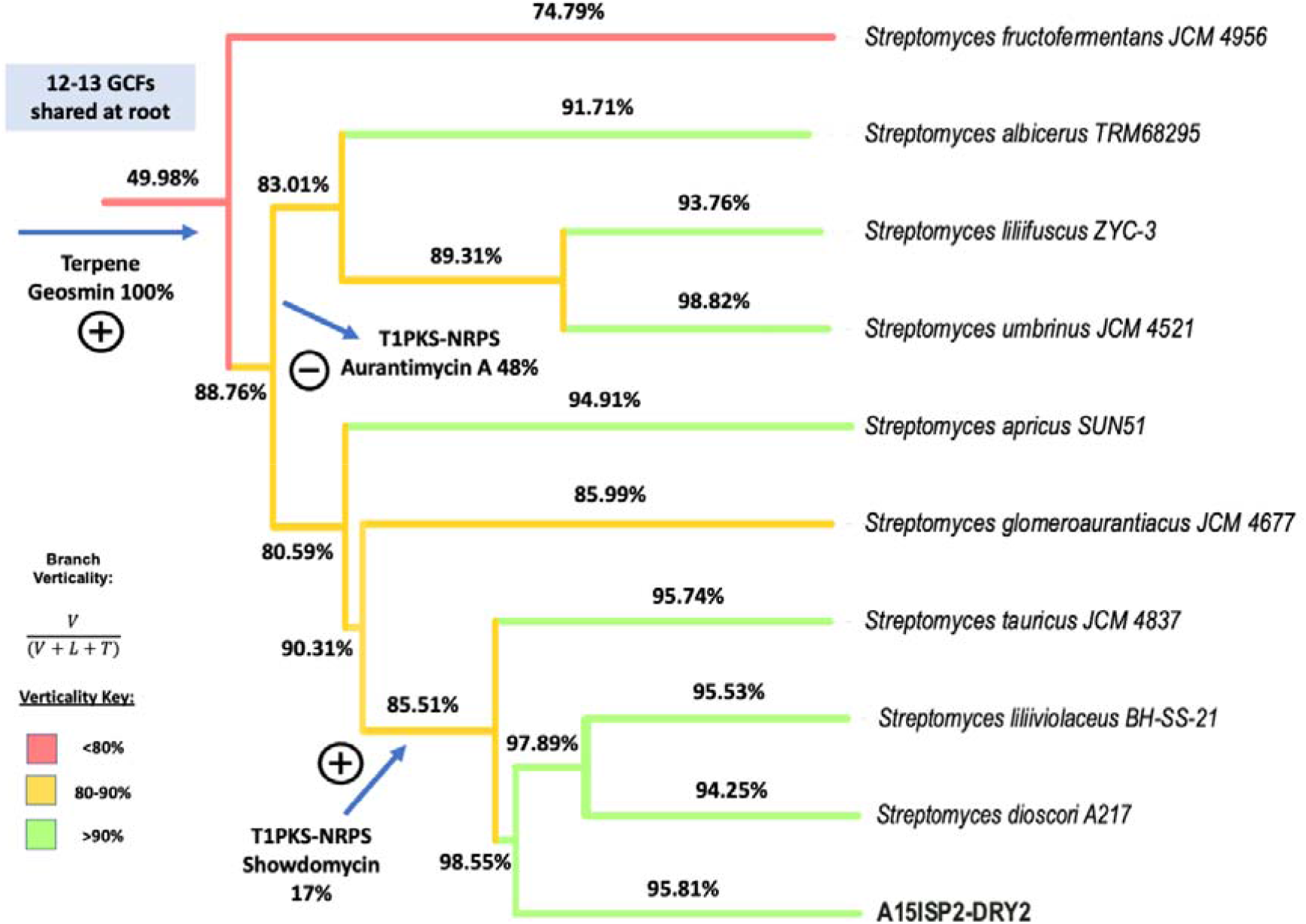
GCF transmission within the S. aurantiacus clade revealed using gene cluster tree-species tree reconciliation. ALE analysis showed the evolution of 28 gene cluster families present in 3 or more strains. These clusters generally evolved vertically, with >90% verticality on most branches of the tree (verticality was measured as the proportion of ancestor-to-descendant vertical transmissions as a proportion of all inferred events). For illustrative purposes, the inferred origin or loss of 3 clusters is illustrated. V = Number of vertical transmissions, L = Number of loss events, T = Number of transfers.

### Antimicrobial activity testing

Following the bioinformatic analysis, we next tested whether this strain produced any antibiotic compounds during laboratory culture. The strain was grown in a liquid culture medium for 7 days and organic extraction of the liquid culture was performed to create a crude metabolite extract. The extract was a bright cherry red in colour. The bioactivity of this metabolite extract was assessed against a panel of clinically relevant pathogenic bacteria (Figure 6; for a full list of strains tested see Materials and Methods). Antibacterial activity was observed against Gram-positive strains including a clinical isolate of vancomycin-resistant and methicillin-resistant *Staphylococcus aureus* designated strain Mu50 (38) and the fusidic acid/rifampicin-resistant *Enterococcus faecalis* JH2-2 (39). The antibiotic activity of the culture extract was somewhat dependent on the culture conditions used; for example, the antibiotic activity was drastically reduced in extracts using mannitol as the carbon source (Figure S3). Further work is now ongoing to determine the extract component responsible for this bioactivity.

**Figure 6.**
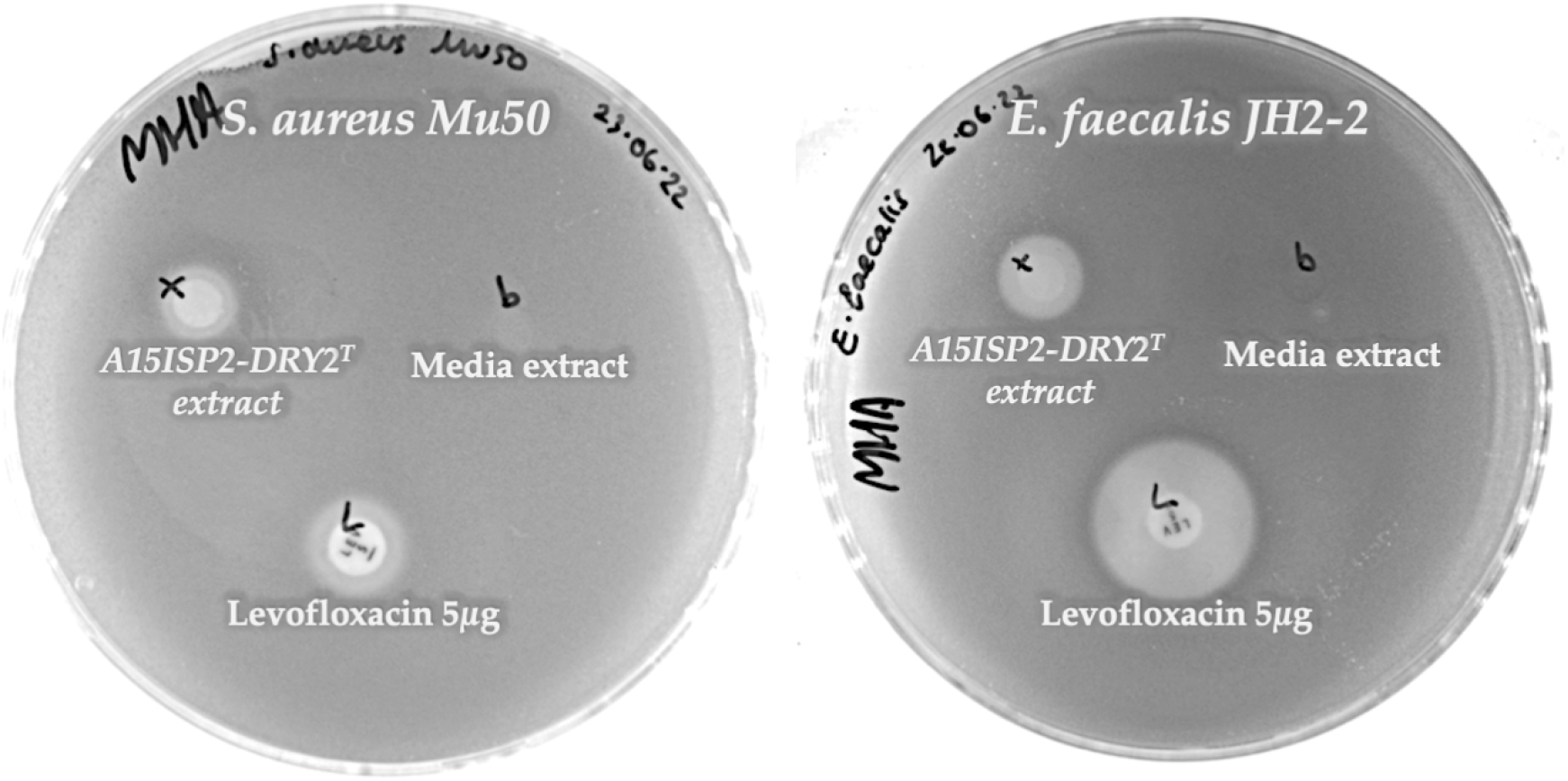
Example of antibiotic activity in crude media extracts of Streptomyces A15T. Clearance zones are visible in agar overlay assays against the pathogens Staphylococcus aureus Mu50 (left) and Enterococcus faecialis JH2-2 (right). The broad-spectrum antibiotic Lexofloxacin is used as a positive control.

## Discussion

The results presented here outline the discovery of a novel species of *Streptomyces* and represent the first *Streptomyces* strain isolated in our wider efforts to isolate deep-sea sponge-associated microbes (26). This study provides evidence that continuing to isolate actinomycetes, even those mined extensively such as *Streptomyces*, can lead to the discovery of taxonomic novelty and biosynthetic diversity. This supports prior large-scale global sequence analyses demonstrating that *Streptomyces*, in particular *Streptomyces*_RG1, not only have the highest number of GCFs among bacteria, but also the highest number of yet uncharacterised and unknown GCFs (24). The current study indicates that continuing to isolate, sequence, and screen novel species within this group could represent an important route to the discovery of new natural products.

Our study suggests that GCF content is shaped both by vertical inheritance and lateral gene transfer; strains share much of their GCF repertoire with close relatives, but also acquire individual GCFs from more distant lineages. Evidence on the extent and rate of LGT within *Streptomyces* is mixed. A phylogenetic study of LGT within Streptomyces suggested that only 10 LGT events occur every million years and that while the transfer of biosynthetic genes was overrepresented in the data, the transfer of entire intact BGCs was relatively rare (40). In contrast, a study by Martinet et al. investigating 18 strains from the same species (*S. lunaelactis)*, found that 54% of BGCs were actually strain-specific (41). This lack of conserved BGCs within a species highlights a limitation of this analysis, as different strains within the same species may have a variable biosynthetic repertoire, driven in part by the acquisition of new clusters from outside the clade. Additional aspects to consider are that fragmented GCFs will decrease the apparent vertical transmission, while the omission of GCFs present in only one or two strains will increase the apparent verticality. So, while it remains clear that phylogenetic distance plays a major role in the biosynthetic repertoire – with A15ISP2-DRY2^T^ sharing most of its GCFs with its closest relatives – exchange of BGCs over larger evolutionary distances is also important, as reflected in the 6 unique GCFs present in A15ISP2-DRY2^T^ but not found elsewhere in the clade (Table S8). This result is consistent with the view that lateral acquisition of BGCs is an important driver of biosynthetic diversity. In the future, a systematic study of *Streptomyces* radiation using these techniques might help to provide a global picture of the relative contributions of vertical descent and LGT to GCF content.

Here we use BiG-SCAPE and ARTS to rapidly assess the biosynthetic novelty of A15ISP2-DRY2^T^ within its clade. Resistance-based mining or target-directed mining — the identification of resistance genes within BGCs — has led to the discovery of several first-in-class antibiotics from *Streptomyces* in recent years (30, 42). ARTS identified that 19 of 34 BGC regions in A15ISP2-DRY2^T^ contained such potential resistance or a duplicated core gene, including two GCFs found exclusively in A15ISP2-DRY2^T^. These clusters represent priority targets for compound isolation or, if not produced under standard laboratory conditions, heterologous expression (43, 44). A15ISP2-DRY2^T^ shared the majority of the identified GCFs with its closest relatives (*S. liliiviolacens* BH-SS-21^T^ and *‘S. dioscori* A217^T^’), despite these strains being isolated from drastically different environments: *S. dioscori* from a yam plant and *S. liliiviolacens* from soil (45, 46). Interestingly, the related strains show distinct bioactivities, with *‘S. dioscori’* A217 having activity solely against Gram-negative *Klebsiella pneumoniae* (46) and *S. liliiviolacens* BH-SS-21^T^ reporting activity against the Gram-negative plant pathogen *Ralstonia solanacearum* (45). This suggests that the metabolomic profile of A15ISP2-DRY2^T^ is distinct from its close relatives and highlights the poorly understood relationship between the genome and metabolome of a strain (47, 48). It is possible that different environmental niches may select for strains with distinct metabolomes rather than distinct BGCs (48). The distinct antibiotic spectrum displayed by our novel deep-sea strain, therefore, warrants further investigation and the characterisation of the specialised metabolite(s) responsible is now a priority. Overall, the *in silico* and *in vitro* results presented here suggest this novel deep-sea *Streptomyces* is an excellent candidate for further antibiotic bioprospecting.

### Description of ‘*Streptomyces ortus* sp. nov.’

The systematic name proposed for the isolated strain A15ISP2-DRY2^T^ is *Streptomyces ortus* sp. nov. The species nomenclature *ortus* (or-tus), sunrise or risen, L. masculine. adj. reflects the deep-sea origins of the original isolate and the red colour that colonies display when grown on ISP2 media. The type strain is A15ISP2-DRY2^T^ (NCIMB 15405^T^ = DSM 113116^T^), isolated from a deep-sea demosponge sponge identified as *Polymastia corticata* from the Gramberg seamount in the Atlantic Ocean (depth 1869m; latitude 15.44530167; longitude: -51.09606). An aerobic, Gram positive, filamentous actinomycetota, that forms branched substrate hyphae and aerial mycelia with spores (∼ 0.82 µm x 1.27 µm) on mannitol soy flour agar. The strain has a high salt tolerance with growth up to 8 % w/v (1.37 M) NaCl. It was successfully cultured at pH 5-12 and at temperatures 4, 15, 20, 28, and 37 ºC. The strain had 6 complete rRNAs with each 16S rRNA gene sequence deposited in GenBank ON356021-ON356026. The genome size of the type strain is 9.27 Mb (GenBank Accession: JAIFZO000000000) and its genomic DNA GC content is 70.83 %.

## Methods

### Sponge identification

The sponge sample was thoroughly rinsed three times in sterile artificial seawater (33.3 g/L, Crystal Sea Marine Mix, Marine Enterprise International). DNA was extracted from ∼0.25 g of sponge tissue in a laminar flow hood using the DNeasy PowerSoil Kit (Qiagen, Hilden, Germany) using the optimized procedure of Marotz *et al*. (49). Sponge taxonomy was based on the mitochondrial cytochrome oxidase subunit I (COI). The COI gene was amplified through PCR using the universal primer pair LCO1490 (GGTCAACAAATCATAAAGATATTGG) and HCO2198 (TAAACTTCAGGGTGACCAAAAAATCA) (50). The reaction comprised 20 µL Platinum™ Hot Start PCR Master Mix (Thermo Fisher Scientific, Waltham, MA, USA), 2 µL of each primer at 10 pmol/µl, 14 µL MilliQ water and 2 µL DNA template. Thermocycler conditions were as described by Yang *et al*. (51): 1-minute denaturation at 94°C; 5 cycles of 94°C for 30□sec, 45°C for 90□sec and 72°C for 1□min; 35 cycles of 94°C for 30□sec, 51°C for 40□sec and 72°C for 1□min; and a final extension step at 72°C for 5□min. The successful amplification of a COI gene fragment of approximately 680 bp was confirmed by agarose gel electrophoresis before the amplicon was purified with the DNA Cleanup Kit (New England Biolabs, Ipswich, MA, USA) and sequenced (Eurofins Genomics, Wolverhampton, UK). The closest relative was determined based on the highest percent identity in a BLASTN search against the NCBI Nucleotide database (52).

### Isolation of A15ISP2-DRY2^T^

The procedures for strain culturing and isolation, and the method of antibiotic screening by soft agar overlay, were performed as previously described (26). To bias strain culturing towards spore-forming bacteria, a dry-stamping technique was adapted from Mincer *et al*. (53). 0.25 g of the sponge was dried at 60ºC for 90 minutes, ground with a sterile pestle, and then stamped onto ISP2 (4.0 g dextrose, 4.0 g yeast extract, 10.0 g malt extract, 33.3 g Crystal Sea Marine Mix, Marine Enterprise International, 15 g agar, 1 L ddH_2_O) agar plates using a sterile plastic bung.

### Cryo-SEM imaging

A15ISP2-DRY2^T^ was streaked onto Mannitol Soya Flour Medium (54), grown for 4 days at 28ºC and a single colony was removed with a scalpel. For scanning electron microscopy (SEM), colonies were mounted on the surface of an aluminium stub with optimal cutting temperature (O.C.T) compound (Agar Scientific) mixed with colloidal graphite as the mounting medium. The stub was plunged into liquid nitrogen slush to cryopreserve the material. Each sample was transferred to the preparation chamber of a Quorum PP3010T cryo-transfer system attached to a JEOL 7900 Field Emission SEM. Sublimation of surface frost was performed at -95°C for 3 minutes before re-cooling then sputter coating with platinum for 2 minutes at 10mA. After coating the sample was transferred to the cryo stage mounted in the SEM chamber held at approximately -140°C. The samples were imaged at 5kV. Average spore dimensions were determined with Fiji (ImageJ v2.3.0) using the UCSB NanoFab plugin Microscope Measurement Tools package.

### Strain Growth Conditions

The standard growth conditions for culturing A15ISP2-DRY2^T^ were on standard ISP2 media (4.0 g dextrose, 4.0 g yeast extract, 10.0 g malt extract, 15 g agar, 1L ddH_2_O) at 28ºC, pH 7.2. To examine the impact of different growth conditions, strains were cultured in triplicate on ISP2 agar over a range of temperatures, salinities, and pH values deviating from the standard culture conditions. Growth temperatures on standard ISP2 were 0ºC, 4°C, 15°C, 20°C, 28°C, and 37°C. Additional NaCl was introduced at concentrations between 0%, 2%, 4%, 6%, 8%, 10%, 12% and 15% w/v. pH was adjusted to final values of 5, 6, 7, 8, 9, 10, 11 and 12 with 2 M HCl or 2 M NaOH. Plates incubated at 0ºC were also supplemented with 3% glycerol to prevent freezing. Gram staining was carried out as previously described (55).

### Analysis of Fatty Acid Cell Wall Composition

Chemotaxonomic analysis was prepared from biomass produced in M.65 medium (L-1) containing 4.0 g glucose, 4.0 g yeast extract, and 10.0 g malt extract (pH 7.2).

Whole-cell sugars and isomers of diaminopimelic acid were diagnosed with standard samples by thin-layer chromatography (TLC) on cellulose plates according to Staneck (56). Polar lipids were extracted from freeze-dried material in chloroform:methanol:0.3% aqueous NaCl, separated by two-dimensional TLC and detected according to Tindall (57). Cellular fatty acids were extracted, methylated and analysed using minor modifications of the methods of Miller (58) and Kuykendall *et al*. (59). The fatty-acid methyl esters were separated by GC and identified using the Sherlock Microbial Identification System (MIDI, Newark, USA). Menaquinones were extracted in hexane, purified by silica-based solid phase extraction, and analysed by reverse phase HPLC-DAD-MS. All analyses were performed by DSMZ services, Leibniz-Institute DSMZ, Braunschweig, Germany.

### Genome assembly

A colony was inoculated from a freshly-streaked ISP2 agar plate into 1mL of liquid media of the same composition. These liquid cultures were incubated in a shaking incubator (180 rpm, 28ºC) for 3-4 days until confluent bacterial growth was achieved. Genome extraction was then performed using the GenElute Bacterial Genomic DNA (gDNA) extraction kit (Sigma-Aldrich, St. Louis, MO, USA) as per the manufacturer’s instructions. Illumina sequencing was performed as a commercial service by Microbes NG (https://microbesng.com/). Briefly, libraries were constructed using the XT Index Kit (Nextera®) and sequenced using HiSeq or NovoSeq platform (Illumina, San Diego, CA, USA) to produce 2 × 250 bp paired-end reads. Trimmomatic (v0.30) was used for adaptor and quality trimming with a sliding window quality cut-off of Q15 (60). Nanopore sequencing was conducted in-house. Extracted genomic DNA was sequenced using a Mk1B R9 MinION flow-cell (Oxford Nanopore Technologies, Oxford, UK) and raw fast5 files were based called using Guppy (v6.3.8). Sequencing files were assembled *de novo* with Unicycler v0.4.6 (61) and the assembly was scaffolded with MeDuSa v1.6 (62) (reference strains listed in Table S8). Alignment of trimmed Illumina reads to the final assemblies used Bowtie2 v2.2.9 (63) and error rate was calculated with Qualimap2 v2.2.2 (64). The contiguity and accuracy of the assembly were assessed respectively with QUAST v5.0.2 (65) and Benchmarking Universal Single-Copy Orthologs (BUSCO) (v5.3.2) (66). Assembly and quality assessment was completed on the Galaxy web platform (https://usegalaxy.eu/) (67).

### Phylogenomics

The final genome assembly file was submitted to the DSMZ Type Strain Genome Server (TYGS v .321; https://tygs.dsmz.de/) and the Genome to Genome Distance Calculator (GGDC v 2.1; https://ggdc.dsmz.de) (27). Phylogenetic trees were visualised using the interactive tree of life (iTOL) V5 (68) with *Micromonospora echinospora* ATCC 15837 selected as an outgroup. Average nucleotide identity (ANI) was calculated on www.usegalaxy.eu using the fastANI algorithm (69) against the closest relatives identified by TYGS. The genome was submitted for annotation to the NCBI Prokaryote Genome Annotation Pipeline (PGAP) (v6.3) (70).

### Biosynthetic Cluster Comparison of the *S. aurantiacus* Clade

BGCs from the *S. aurantiacus* clade and the isolated strain was identified using antiSMASH 6.1 (19). Where multiple assemblies were available, the highest-quality genome for each species in GenBank was chosen (Table S8). To enable cluster comparison, detected BGCs were grouped into Gene Cluster Families (GCF) using BiG-SCAPE v1.1.0 (21). BiG-SCAPE was run with the *--mix* flag and the clustering distance parameter was tested at 0.3, 0.35, 0.4, 0.45 and 0.5. 0.35 was chosen, as this represented the lowest cut-off that grouped the geosmin BGC into a single GCF (71). The resulting network map output was loaded into Cytoscape v3.9.1 (72). The *--mibig* flag was used to identify closely related clusters from The Minimum Information about a Biosynthetic Gene cluster (MiBIG 2.0) database (31). For each GCF, aligned amino acid sequences produced by BiG-SCAPE were used as input for IQTREE2 (73) to create 1000 bootstrapped GCF trees using the best-fitting substitution model as selected by the Bayesian Information Criterion. Bootstrapped trees were used as input for ALEobserve and then reconciled against the rooted species tree using ALEml_undated (ALE v1.0) (37). The verticality of each branch was then calculated as the inferred number of ancestor-to-descendant vertical transmissions as a proportion of all inferred events.

### Bioactivity Testing

A cube of mycelium containing agar was used to inoculate a starter culture of 50mL liquid ISP2 without dextrose in 250 mL baffled Erlenmeyer flasks. After 24 hours of growth at 28°C with agitation at 180 rpm, 10 mL of the starter culture was used to inoculate 100 mL of ISP2 media in 250 mL baffled Erlenmeyer flasks and grown for 7 days (28°C, 180 rpm). The total culture was then extracted through rigorous shaking with an equal volume of ethyl acetate (Sigma-Aldrich, St. Louis, MO, USA). The organic layer was removed and dried over anhydrous MgSO_4_ (Sigma-Aldrich, St. Louis, MO, USA) before being evaporated under vacuum. The dried crude extract was resuspended in 2 mL methanol for bioactivity testing.

Bioactivity testing employed the following ESKAPE pathogens; *Staphylococcus aureus* Mu50, *Klebsiella pneumoniae* NCTC 5055, *Acinetobacter baumannii* ATCC 19606, *Pseudomonas aeruginosa* PAO1, *Escherichia coli* BW55113 and *Enterococcus faecalis* UB591 (JH2-2). All strains were grown in Muller-Hinton broth at 37°C for 16 h with shaking at 180 rpm. Cultures were diluted to OD_600_ = 0.01 in warm, molten Muller-Hinton agar (0.75 % agar), and 10 mL was poured over a plate of solid Muller-Hinton agar and allowed to set. The crude extract of A15ISP2-DRY2^T^ (25 µL) was dried onto sterile disks of filter paper, which were placed onto the surface of the set agar. ISP2 media extracts were used as a negative control and disks containing Levofloxacin (5 µg) were used as positive controls. The plates were incubated at 37°C, for 16 h, at which point zones of inhibition could be observed.

## Data Availability

The partial COI gene was deposited in GenBank with the accession OP036683. In total A15ISP2-DRY2 had six 16S rRNA genes. Each was submitted to the NCBI 16S rRNA database with the accession numbers ON356021-ON356026. This Whole Genome Shotgun project has been deposited at DDBJ/ENA/GenBank under the accession JAIFZO000000000. The version described in this paper is version JAIFZO010000000.

## Contributions

Conceptualization: SEW, PC, PRR

Methodology: SEW, TAW, JM

Investigation: SEW, CB, EB, JS, JM

Visualization: SEW

Funding acquisition: PC, PRR

Supervision: PC, PRR

Formal analysis: SEW, TAW

Writing – original draft: SEW, PC

Writing – review & editing: SEW, PC, CB, PRR, EB, TAW

Conflicts of Interest: Authors declare no potential conflicts of interest

## Funding and acknowledgements

SEW is supported by the Bristol Centre for Engineering Biology (BrisEngBio) under UKRI Biotechnology and Biological Sciences (BBSRC) award BB/W013959/1. Further grants that have supported this work include BBSRC grants BB/T001968/1 and BB/M025624/1, and the Medical Research Foundation grant MRF-131-0005-RG-RACE-C0853. EB was supported by the Wellcome Trust ISSF and Elizabeth Blackwell Institute clinical primer scheme. The TROPICS research cruise (expedition JC094) was funded by the European Research Council via the ERC Consolidator Grant agreement number 278705. TAW is supported by a Royal Society University Research Fellowship (URF\R\201024).

The authors gratefully acknowledge the Material and Chemical Characterisation Facility (MC^2^) at the University of Bath (doi.org/10.15125/mx6j-3r54) for technical support and assistance in obtaining Cryo-SEM images for the paper. The authors also thank Laura Robinson and Kate Hendry for sample collection during expedition JC094.

