## Supplementary Materials for "Discovery and biosynthetic assessment of *Streptomyces ortus* sp nov. isolated from a deep-sea sponge"

Supplementary material:

Table S1. Top 10 BLASTN (2.13.0+) hits for the deep-sea sponge 700bp COI gene sequence. NCBI accession: OP036683

| **Description** | **Scientific Name** | **Query Cover** | **% Identity** |
| --- | --- | --- | --- |
| [Polymastia corticata mitochondrial partial COI gene for cytochrome oxidase subunit 1](https://blast.ncbi.nlm.nih.gov/Blast.cgi#alnHdr_563404438) | [*Polymastia corticata*](https://www.ncbi.nlm.nih.gov/Taxonomy/Browser/wwwtax.cgi?id=475993) | 92% | **100%** |
| [Polymastia littoralis mitochondrion, complete genome](https://blast.ncbi.nlm.nih.gov/Blast.cgi#alnHdr_594595143) | [*Polymastia littoralis*](https://www.ncbi.nlm.nih.gov/Taxonomy/Browser/wwwtax.cgi?id=1473587) | 99% | **95.76%** |
| [Polymastia atlantica voucher TS2947 cytochrome c oxidase subunit I (COX1) gene](https://blast.ncbi.nlm.nih.gov/Blast.cgi#alnHdr_1843765328) | [*Polymastia atlantica*](https://www.ncbi.nlm.nih.gov/Taxonomy/Browser/wwwtax.cgi?id=2737414) | 94% | **96.31%** |
| [Polymastia atlantica voucher TS2938 cytochrome c oxidase subunit I (COX1) gene](https://blast.ncbi.nlm.nih.gov/Blast.cgi#alnHdr_1843765326) | [*Polymastia atlantica*](https://www.ncbi.nlm.nih.gov/Taxonomy/Browser/wwwtax.cgi?id=2737414) | 94% | **96.31%** |
| [Polymastia sp. 2 PRT-2020 voucher TS3976 cytochrome c oxidase subunit I (COX1) gene](https://blast.ncbi.nlm.nih.gov/Blast.cgi#alnHdr_1843765344) | [*Polymastia sp. 2 PRT-2020*](https://www.ncbi.nlm.nih.gov/Taxonomy/Browser/wwwtax.cgi?id=2737348) | 94% | **96.02%** |
| [Sphaerotylus strobilis voucher TS3628 cytochrome c oxidase subunit I (COX1) gene](https://blast.ncbi.nlm.nih.gov/Blast.cgi#alnHdr_1843765338) | [*Sphaerotylus strobilis*](https://www.ncbi.nlm.nih.gov/Taxonomy/Browser/wwwtax.cgi?id=2737413) | 94% | **95.87%** |
| [Sphaerotylus strobilis voucher TS4700 cytochrome c oxidase subunit I (COX1) gene](https://blast.ncbi.nlm.nih.gov/Blast.cgi#alnHdr_1843765336) | [*Sphaerotylus strobilis*](https://www.ncbi.nlm.nih.gov/Taxonomy/Browser/wwwtax.cgi?id=2737413) | 94% | **95.87%** |
| [Sphaerotylus strobilis voucher TS4699 cytochrome c oxidase subunit I (COX1) gene](https://blast.ncbi.nlm.nih.gov/Blast.cgi#alnHdr_1843765334) | [*Sphaerotylus strobilis*](https://www.ncbi.nlm.nih.gov/Taxonomy/Browser/wwwtax.cgi?id=2737413) | 94% | **95.87%** |
| [Sphaerotylus strobilis voucher TS4697 cytochrome c oxidase subunit I (COX1) gene](https://blast.ncbi.nlm.nih.gov/Blast.cgi#alnHdr_1843765332) | [*Sphaerotylus strobilis*](https://www.ncbi.nlm.nih.gov/Taxonomy/Browser/wwwtax.cgi?id=2737413) | 94% | **95.87%** |
| [Sphaerotylus strobilis voucher TS2685 cytochrome c oxidase subunit I (COX1) gene](https://blast.ncbi.nlm.nih.gov/Blast.cgi#alnHdr_1843765324) | [*Sphaerotylus strobilis*](https://www.ncbi.nlm.nih.gov/Taxonomy/Browser/wwwtax.cgi?id=2737413) | 94% | **95.87%** |

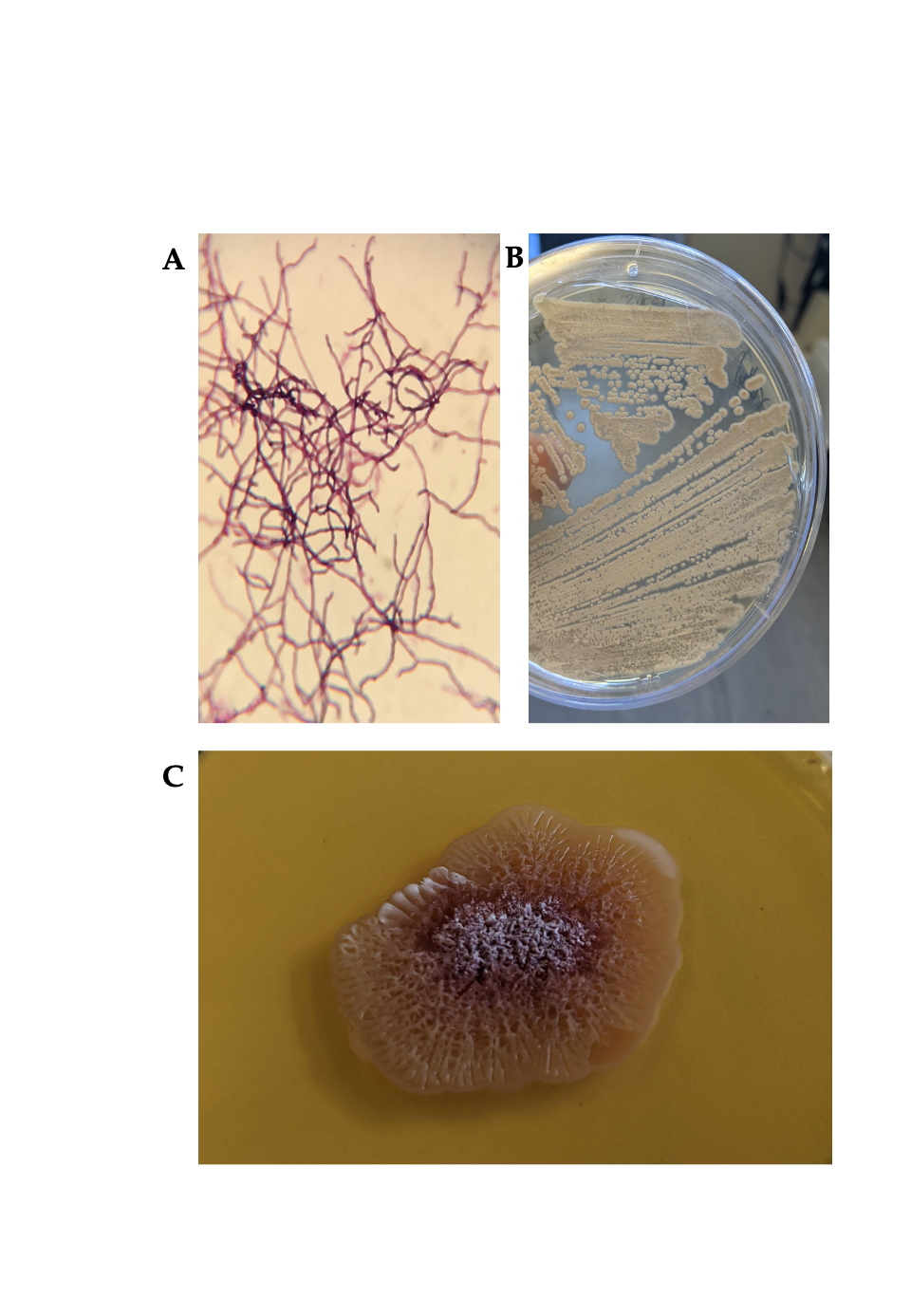

Figure S1. Images of microbial isolate A15ISP2-DRY2 ^T^. (A) The strain was Gram positive and showed filamentous structures when viewed under a light microscope. (B) Colonies were initially cream, but over time changed to orange and finally burgundy in colour (C) The strain had the ability to form white spores on ISP2 agar

Table S2. Summary statistics for the genome assembly of strain Streptomyces A15. Assembled using Unicycler and scaffolded with MeDuSa, assembly metrics assessed with QUAST (contigs >= 500bp), Bowtie2 and Qualimap.

| **Assembly metric** | **Assembly** |
| --- | --- |
| **Number of contigs** | 9 |
| **Number of scaffolds** | 4 |
| **Total length (Mb)** | 9.29 |
| **Largest contig (Mb)** | 5.23 |
| **Largest scaffold (Mb)** | 8.61 |
| **Coverage** | 77.6391 |
| **N50 (Mb)** | 8.61 |
| **L50** | 1 |
| **Mapped reads** | 98.72% |
| **Error rate** | 0.8% |
| **GC content** | 70.83% |

Table S3. NCBI Prokaroyate genome annotation pipeline results for genome

| Annotation Provider | NCBI |
| --- | --- |
| Annotation Date | 10/21/2022 13:51:25 |
| Annotation Pipeline | NCBI (PGAP) |
| Annotation Method | Best-placed reference protein set; GeneMarkS-2+ |
| Annotation Software revision | 6.3 |
| Features Annotated | Gene; CDS; rRNA; tRNA; ncRNA; repeat_region |
| Genes (total) | 8,130 |
| CDSs (total) | 8,043 |
| Genes (coding) | 7,794 |
| CDSs (with protein) | 7,794 |
| Genes (RNA) | 87 |
| rRNAs | 6, 6, 6 (5S, 16S, 23S) |
| complete rRNAs | 6, 6, 6 (5S, 16S, 23S) |
| tRNAs | 66 |
| ncRNAs | 3 |
| Pseudo Genes (total) | 259 |
| CDSs (without protein) | 259 |
| Pseudo Genes (ambiguous residues) | 0 of 259 |
| Pseudo Genes (frameshifted) | 80 of 259 |
| Pseudo Genes (incomplete) | 193 of 259 |
| Pseudo Genes (internal stop) | 19 of 259 |
| Pseudo Genes (multiple problems) | 40 of 259 |
| CRISPR Arrays | 1 |

Table S4. Table of single-copy orthologous genes present in the Streptomyces A15 genome assembly as expected for a genome from the order streptomycetales (lineage dataset: streptomycetales_odb10).

| **Assembler** | **Complete**  **[Single copy/Duplicated]** | **Fragmentated** | **Missing** | **Number of BUSCO groups searched** |
| --- | --- | --- | --- | --- |
| **Unicycler** | 99.7% [99.4%/0.3%] | 0.1% | 0.2% | 1579 |

Table S5. GGDC formula 2 (d4) dDDH (TGYS)and ANI values for 10 closely related strains to A15ISP2-DRY2^T^ and M. echinospora ATCC 15837 used as an outgroup

| **Strain** | **dDDH (d4, in %)** | **C.I. (d4, in %)** | **G+C % difference** | **FASTANI (%)** |
| --- | --- | --- | --- | --- |
| *Streptomyces liliiviolaceus* BH-SS-21 | 45.8 | [43.3 - 48.4] | 0.04 | 93.3053 |
| *Streptomyces dioscori* A217 | 45.1 | [42.6 - 47.7] | 0.11 | 93.0804 |
| *Streptomyces tauricus* JCM 4837 | 43.7 | [41.2 - 46.3] | 0 | 92.7929 |
| *Streptomyces glomeroaurantiacus* JCM 4677 | 35.2 | [32.8 - 37.7] | 0.54 | 89.7836 |
| *Streptomyces apricus* SUN51 | 34.7 | [32.3 - 37.2] | 1.27 | 89.4996 |
| *Streptomyces liliifuscus* ZYC-3 | 31.6 | [29.2 - 34.1] | 0.61 | 87.8961 |
| *Streptomyces umbrinus* JCM 4521 | 31.6 | [29.2 - 34.1] | 0.66 | 87.8979 |
| *Streptomyces albicerus* TRM68295 | 31.5 | [29.1 - 34.0] | 0.81 | 87.5189 |
| *Streptomyces ederensis* JCM 4958 | 31.5 | [29.1 - 34.0] | 0.46 | 86.239 |
| *Streptomyces fructofermentans* JCM 4956 | 29.2 | [26.8 - 31.7] | 1.58 | 83.6057 |
| *Streptomyces stelliscabiei* DSM 41803 | 25.1 | [22.8 - 27.6] | 0.27 | 83.554 |
| *Streptomyces caniscabiei* NE06-02D | 25 | [22.7 - 27.5] | 0.45 | 83.5369 |
| *Micromonospora echinospora* ATCC 15837 | 18.9 | [16.7 - 21.2] | 1.54 | 75.9895 |

Table S6. antiSMASH results with closest ClusterBlast hit 6.1.1 and antiSMASH db 3.0

| Region | Type | ClusterBlast Hit antiSMASHdb | Most Similar Known Cluster | Similarity |
| --- | --- | --- | --- | --- |
| 1 | NRPS, T3PKS | Streptomyces dioscori strain A217 (100%) | Herboxidiene | 10% |
| 2 | Terpene | Streptomyces dioscori strain A217 (100%) | 2-methylisoborneol | 100% |
| 3 | NRPS | Streptomyces albireticuli strain MDJK11 (23%) | Foxicins A-D | 29% |
| 4 | Siderophore | Streptomyces dioscori strain A217 (87%) | *No similar cluster* |  |
| 5 | NAPAA | Streptomyces dioscori strain A217 (58%) | Rapamycin | 17% |
| 6 | Ecotine | Streptomyces dioscori strain A217 (100%) | Ecotine | 100% |
| 7 | Terpene | Streptomyces dioscori strain A217 (85%) | Albaflavenone | 100% |
| 8 | PKS-like, T1PKS | Streptomyces coelicolor A3(2) CFB_NBC_0001 (70%) | Arsono-polyketide | 91% |
| 9 | T3PKS | Streptomyces sp. CS131 (30%) | Alkylresorcinol | 66% |
| 10 | NRPS, T3PKS, terpene | Streptomyces aquilus strain GGCR-6 (37%) | Feglymycin | 68% |
| 11 | T2PKS, Ladderane | Streptomyces sp. 3214.6 (53%) | Simocyclinone D8 | 40% |
| 12 | Terpene | Streptomyces dioscori strain A217 (40%) | Isorenieratene | 100% |
| 13 | NRPS | Streptomyces dioscori strain A217 (36%) | Borrelidin | 4% |
| 14 | NRPS | Streptomyces dioscori strain A217 (89%) | Rimosamide | 21% |
| 15 | NRPS | Streptomyces dioscori strain A217 (56%) | Diisonitrile antibiotic SF2768 | 55% |
| 16 | Lanthipeptide- class-iii, RiPP-like | Streptomyces sp. S1A1-3 (100%) | Informatipeptin | 42% |
| 17 | T1PKS, terpene | Streptomyces sp. VN1 (81%) | Oxalomycin B | 9% |
| 18 | Terpene | Streptomyces dioscori strain A217 (60%) | Herboxidiene | 4% |
| 19 | NRPS, NRPS-like, T1PKS, other, terpene | Streptomyces griseochromogenes strain ATCC 145 (49%) | Aurantimycin A | 48% |
| 20 | Terpene | Streptomyces dioscori strain A217 (100%) | Hopene | 92% |
| 21 | NRPS-like, PKS-like, T1PKS, ecotine | Streptomyces dioscori strain A217 (65%) | Showdomycin | 17% |
| 22 | Siderophore | Streptomyces dioscori strain A217 (100%) | Grincamycin | 8% |
| 23 | NAPAA | Streptomyces bottropensis ATCC 25435 (63%) | Stenothricin | 13% |
| 24 | Terpene | Streptomyces dioscori strain A217 (63%) | Geosmin | 100% |
| 25 | RiPP-like | Streptomyces sp. YIM 130001 DSC45 s06 (80%) | *No similar cluster* |  |
| 26 | NRPS, NRPS-like, betalactone | Amycolatopsis alba DSM 44262 (42%) | Vazabitide A | 23% |
| 27 | Lanthipeptide class iv | Streptomyces yanglinensis strain CGMCC 4.2023 (25%) | *No similar cluster* |  |
| 28 | Siderphore | Streptomyces dioscori strain A217 (100%) | *No similar cluster* |  |
| 29 | PKS-like, RRE-containing, T2PKS | Streptomyces dioscori strain A217 (84%) | Cinerubin B | 100% |
| 30 | Melanin | Streptomyces dioscori strain A217 (75%) | Melanin | 60% |
| 31 | Siderophore | Streptomyces dioscori strain A217 (100%) | Desferrioxamin B/E | 83% |
| 32 | RiPP-like | Streptomyces geranii strain A301 (88%) | *No similar cluster* |  |
| 33 | Nucleoside | Streptomyces dioscori strain A217 (66%) | *No similar cluster* |  |
| 34 | NRPS | Streptomyces sp. 150FB (15%) | Lysolipin I | 4% |

Table S7. Type strains of Streptomyces aurantiacus clade used in this study. NCBI GenBank accession numbers listed, BGC identified with antiSMASH 6.0, contigs listed not including scaffolds

| *Streptomyces* species | Assembly Accession | BGCs | Contigs | Assembly size (mb) |
| --- | --- | --- | --- | --- |
| *Streptomyces umbrinus* JCM 4521^T^ | *BMUM00000000* | 41 | 71 | 11.86 |
| *Streptomyces glomeroaurantiacus* JCM 4677 ^T^ | AP023440 | 28 | 1 | 9.43 |
| ‘*Streptomyces dioscori’* A217 | *PYBJ00000000* | 43 | 57 | 10.24 |
| *Streptomyces tauricus* JCM 4837^T^ | *BMVY00000000* | 39 | 107 | 11.04 |
| *Streptomyces liliifuscus* sp. nov. ZYC-3^T^ | CP066831 | 30 | 2 | 11.03 |
| *Streptomyces liliiviolaceus* sp. nov. BH-SS-21^T^ | JAGPYQ000000000 | 39 | 4 | 11.13 |
| *Streptomyces fructofermentans* JCM 4956^T^ | BMWD00000000 | 26 | 116 | 8.87 |
| *Streptomyces albicerus* TRM68295 ^T^ | VWMY00000000 | 48 | 294 | 11.97 |
| *Streptomyces apricus* SUN51^T^ | VDFC00000000 | 43 | 692 | 8.81 |
| *Streptomyces ortus* sp. nov. A15 ^T^ | JAIFZO000000000 | 34 | 9 | 9.29 |

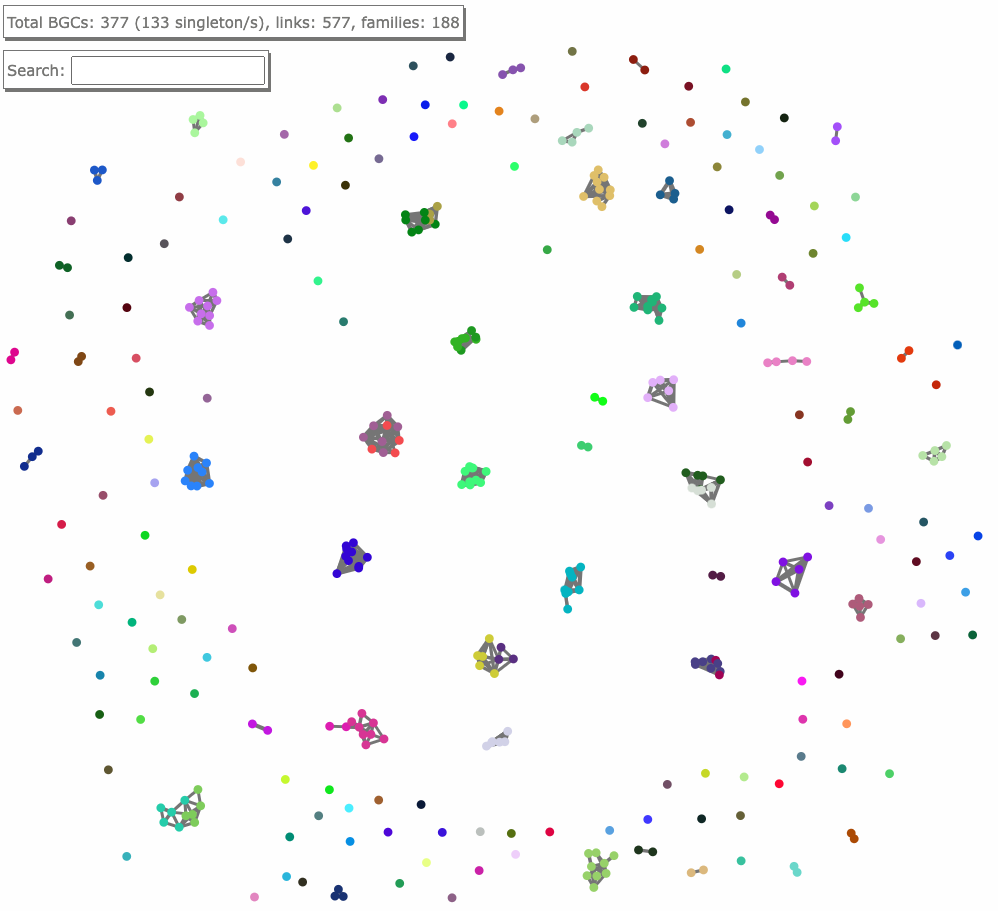

Figure S2. Full BiG-SCAPE (v1.1) GCF family network from the S. auranticus clade including singletons. GCF clustering cutoff 0.35. Direct screen capture from index.html file

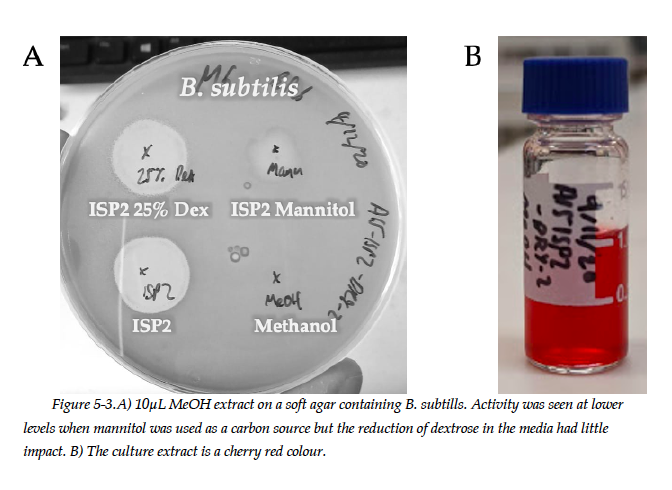

Figure S3. A) Bioactivity of 10µL crude extract on soft agar containing B. subtilis. The activity was reduced when the strain was grown with mannitol instead of dextrose but activity was unaffected if dextrose concentration was reduced to 1g/L (25% of standard). B) The culture extract was a cherry red colour

Table S8: BiG-FAM database information of GCF singletons found in S. ortus. Total does not include putative members.

| **BGC Class** | **Most similar known cluster** | **BiG-FAM family** | **Comment on taxonomy of BiG-FAM family** | **Core member?** |
| --- | --- | --- | --- | --- |
| Siderophore | No similar cluster | GCF_01140 | Staphlococcus 62.9%. Streptomyces 2.8% (6391 total) | TRUE |
| NRPS | NRPS Foxicins A-D 29% | GCF_09952 | Rhodococcus with 7 found in streptomyces (22 total) | FALSE |
| Terpene | Isorenieratene 100% | GCF_00998 | 72% Streptomyces (211 total) | FALSE |
| T2PKS, Ladderane | Simocyclinone D8 40% | GCF_03444 | Rare only 2 BGCs in GCF. All streptomyces | FALSE |
| NRPS | Lysolipin I 4% | GCF_00012 | Mainly found in Pseudomonas – (25972 total) | FALSE |
| NRPS,T3PKS, Terpene | Feglymycin 73% | GCF_00057 | Nonomuraea 2 core members 15 putative (2 total) | FALSE |
